# Validation of ferroptosis in canine cancer cells to enable comparative oncology and translational medicine

**DOI:** 10.1101/2024.04.28.591561

**Authors:** Priya Chatterji, Gang Xing, Laura Furst, Krishna Dave, Qiong Zhou, Daniel V. LaBarbera, Douglas H. Thamm, John K. Eaton, Mathias J. Wawer, Vasanthi S. Viswanathan

**Affiliations:** Kojin Therapeutics, 451 D Street, Suite 502, Boston, MA 02210; The CU Anschutz Center for Drug Discovery, Skaggs School of Pharmacy and Pharmaceutical Sciences, Department of Pharmaceutical Sciences, University of Colorado Anschutz Medical Campus, 12850 E. Montview Blvd, Aurora, CO 80045; Flint Animal Cancer Center, Colorado State University, 300 West Drake Road, Fort Collins, CO 80523

## Abstract

Ferroptosis is a cell death mechanism that has attracted significant attention as a potential basis for the development of new cancer therapies. Validation of ferroptosis biology in species commonly used in translation and pre-clinical development is a necessary foundation for enabling the advancement of such ferroptosis modulating drugs. Here, we demonstrate that canine cancer cells exhibit sensitivity to a wide range of ferroptosis-inducing perturbations in a manner indistinguishable from human cancer cells, and recapitulate characteristic patterns of ferroptotic response across tumor types seen in the human setting. The foundation provided herein establishes the dog as a relevant efficacy and toxicology model for ferroptosis and creates new opportunities to leverage the canine comparative oncology paradigm to accelerate the development of ferroptosis-inducing drugs for human cancer patients.

## Introduction

Ferroptosis is a form of regulated necrosis mediated by the iron-dependent generation of membrane phospholipid peroxides and subsequent loss of membrane integrity.^1^ Susceptibility to ferroptosis has been shown to be a unique vulnerability of cancer cells that exhibit either *de novo* or acquired resistance to apoptosis-inducing therapies.^2,3^ As a newly elucidated biological pathway that has not previously been targeted therapeutically, the ferroptosis field stands to benefit greatly from tools and approaches that can enable demonstration of biological activity, proof of mechanism and evidence of safety in the oncology clinical setting as early as possible in the drug discovery and development process.

The setting of canine cancers represents a powerful resource in this regard: a combination of access to patients with tumors and a clinical trial infrastructure that is uniquely well-suited to enable translational work.^4–6^ Canine cancers are spontaneously developing tumors that can faithfully recapitulate many aspects of the human disease including molecular pathogenesis, histopathology, clinical presentation and response to therapy.^4–6^ Of particular significance to ferroptosis, the incidence of mesenchymal tumors is far greater in the canine setting than among humans: while sarcomas are relatively rare in humans, roughly 15% of canine cancers are sarcomas.^7,8^ This increased incidence translates directly into larger cohorts of patients with ferroptosis-relevant tumors as well as greater access to tissue samples, cell lines and other research tools representing ferroptosis-susceptible cancer states.^7,8^

In addition to biological rationale, clinical trials in the veterinary oncologic setting afford a number of significant advantages for investigation of experimental therapeutics. These features include ability to administer investigational agents in the pre-IND stage, high rates of clinical trial participation and patient compliance, rapid accrual of patients and accelerated timelines for trial completion, and high rates of consent for repeat biospecimen collection to enable in-depth PK-PD and biomarker studies aimed at demonstrating proof of mechanism.^4–6^ The integration of canine trials into the clinical development path has played a key role in the advancement and approval of several important oncology drugs.^9–14^ To enable such approaches for ferroptosis-inducing drugs, we sought to test whether ferroptosis sensitivity exists in canine cancer cells and whether the hallmarks of ferroptosis sensitivity seen across human cancer types is conserved in the canine context.

## Results

We gained access to 31 previously established canine cancer cell lines^15^ (Supplementary Taable 1) and profiled their sensitivity to a panel of 14 compounds covering a diversity of ferroptosis-inducing and other cell killing mechanisms (Figure 1). Compounds were profiled in duplicate across nine dose points, with and without ferrostatin-1,^1,16,17^ a potent and specific inhibitor of ferroptosis. For subsequent analyses, cellular response to treatment was summarized as the area under the dose-response curve (AUC) for each compound–cell line pair. In total, we collected 868 dose–response curves and over 15,600 individual data points.

**Figure 1.**
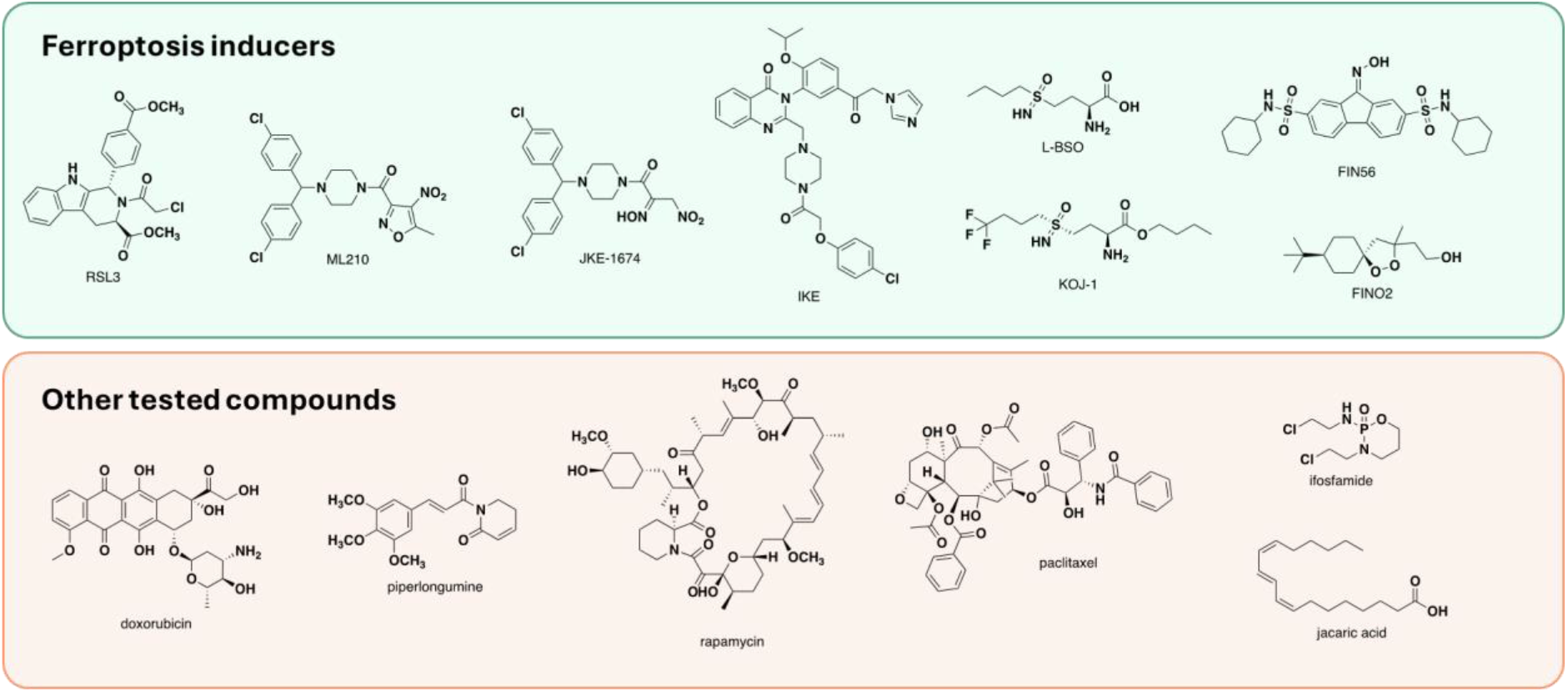
Chemical structures of the 14 compounds tested against the canine cell line panel.

We found that canine cell lines exhibited strong sensitivity to all major classes of ferroptosis inducing perturbations tested (Figure 2A), including GPX4 modulators (RSL3, ML210, JKE-1674, and FIN56),^18–20^ a cystine transport inhibitor (IKE),^21,22^ inhibitors of glutathione biosynthesis (L-BSO and KOJ-1),^23,24^ and pro-oxidants (FINO2 and jacaric acid).^20,25^ The ability of ferrostatin-1 cotreatment to rescue these cell killing responses confirms the ferroptotic nature of cell death of canine cancer cells in response to ferroptosis-inducing perturbations (Figure 2B). Contrary to literature reports, addition of the conjugated polyunsaturated fatty acid (PUFA) jacaric acid^26^ did not have significant effects on cell viability.

**Figure 2.**
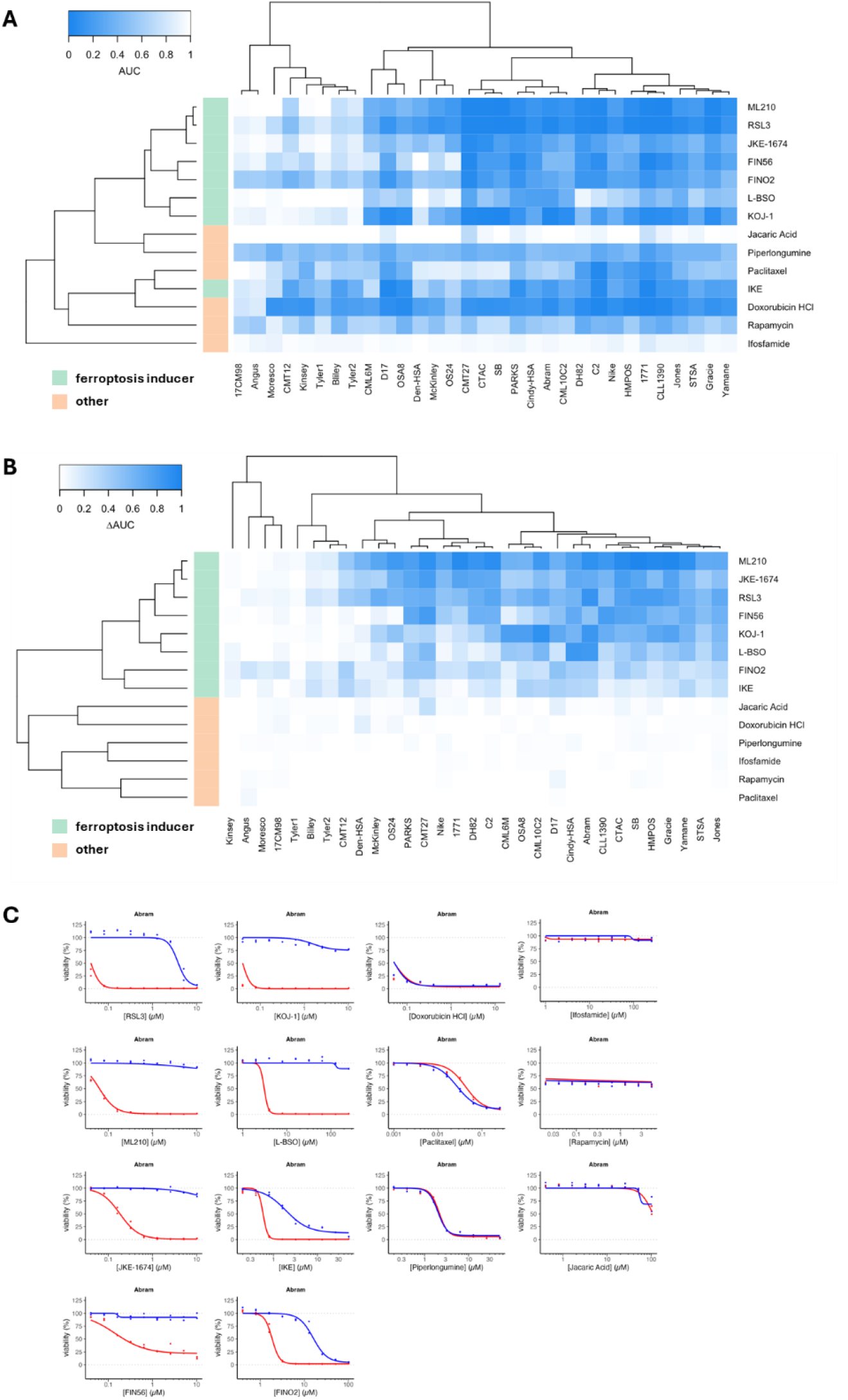
Ferroptosis inducers induce similar cell-killing patterns across canine cancer cell lines, and cell death is rescuable by ferrostatin-1. (A) Heatmap of dose–response AUC values (lower = more sensitive) for all compound–cell line pairs tested. Rows and columns were hierarchically clustered using complete linkage on correlation distances (1-Pearson correlation coefficient). (B) Heatmap of ∆AUC values (higher = better rescue, 0 = no rescue), comparing the difference in AUC between treatment with and without ferrostatin-1. In sensitive cell lines, cell viability effects for all ferroptosis inducers (green) are rescued by ferrostatin-1, while ferrostatin-1 treatment has little effect on other compounds (red). (C) Dose–response curves for all tested compounds, exemplified via cell line Abram.

Like the human context, canine cell lines displayed preferential ferroptosis sensitivity according to their tumor types of origin.^2^ When clustered based on AUC values for ferroptosis inducers, cell lines separated into two main groups that correspond to ferroptosis-sensitive and insensitive cell lines, respectively. Epithelial cancers (carcinomas) were enriched in the ferroptosis-insensitive cluster, while sarcomas, undifferentiated melanomas and hematological malignancies were enriched for sensitivity to ferroptosis (Figure 3A). Control lethal agents did not show this cell state-specific cell killing (Figure 3B).

**Figure 3.**
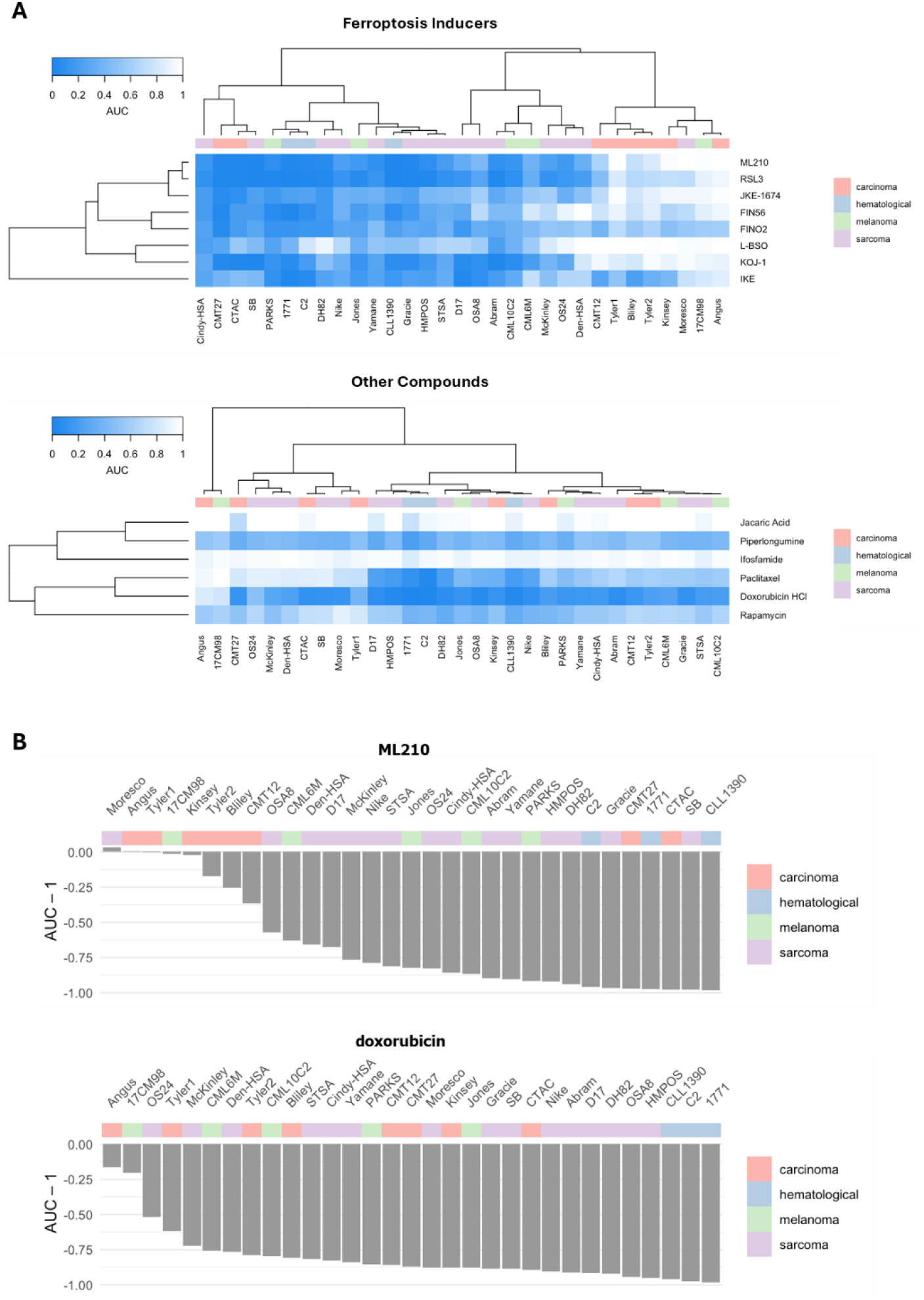
Canine cancer cell lines cluster by tumor type based on sensitivity to ferroptosis inducers. (A) Heatmaps of cell viability AUC values (lower = more sensitive) for ferroptosis inducers (top) and other compounds (bottom). Enrichment of epithelial cancer types in the insensitive cluster (top right) is only observed for ferroptosis inducers. (B) Bar charts indicating the degree of sensitivity (AUC-1, lower value = more sensitive) for ML210 (GPX4 inhibitor; top) and doxorubicin (topoisomerase inhibitor; bottom). Rank ordering cell lines by sensitivity indicates selectivity for killing non-epithelial cells for ML210 but not doxorubicin.

To further characterize the tested canine cell lines, we collected untargeted lipidomic data for each line in duplicate. Omics characterizations of cancer cell lines and integration with viability data in efforts such as the Cancer Cell Line Encyclopedia^27,28^ and the Cancer Dependency Map (www.depmap.org) have yielded powerful resources to explore causal and predictive biomarkers of response to genetic and small molecule perturbations. Our data facilitate a similar approach for the canine lines included in this study.

Lipidomics data covered over 750 individual species falling into 18 main lipid classes (Figure 4A). Total concentrations varied widely between individual cell lines (Figure 4B), as did relative lipid composition (Figure 4C, Supporting Figure 1). Likewise, the fraction of cellular polyunsaturated fatty acids (PUFAs), previously reported as a contributor to ferroptosis sensitivity,^29^ varied from 20% to almost 70% (Figure 4D). Thus, the panel of tested canine cell lines represented a wide spectrum of cellular lipid composition.

**Figure 4.**
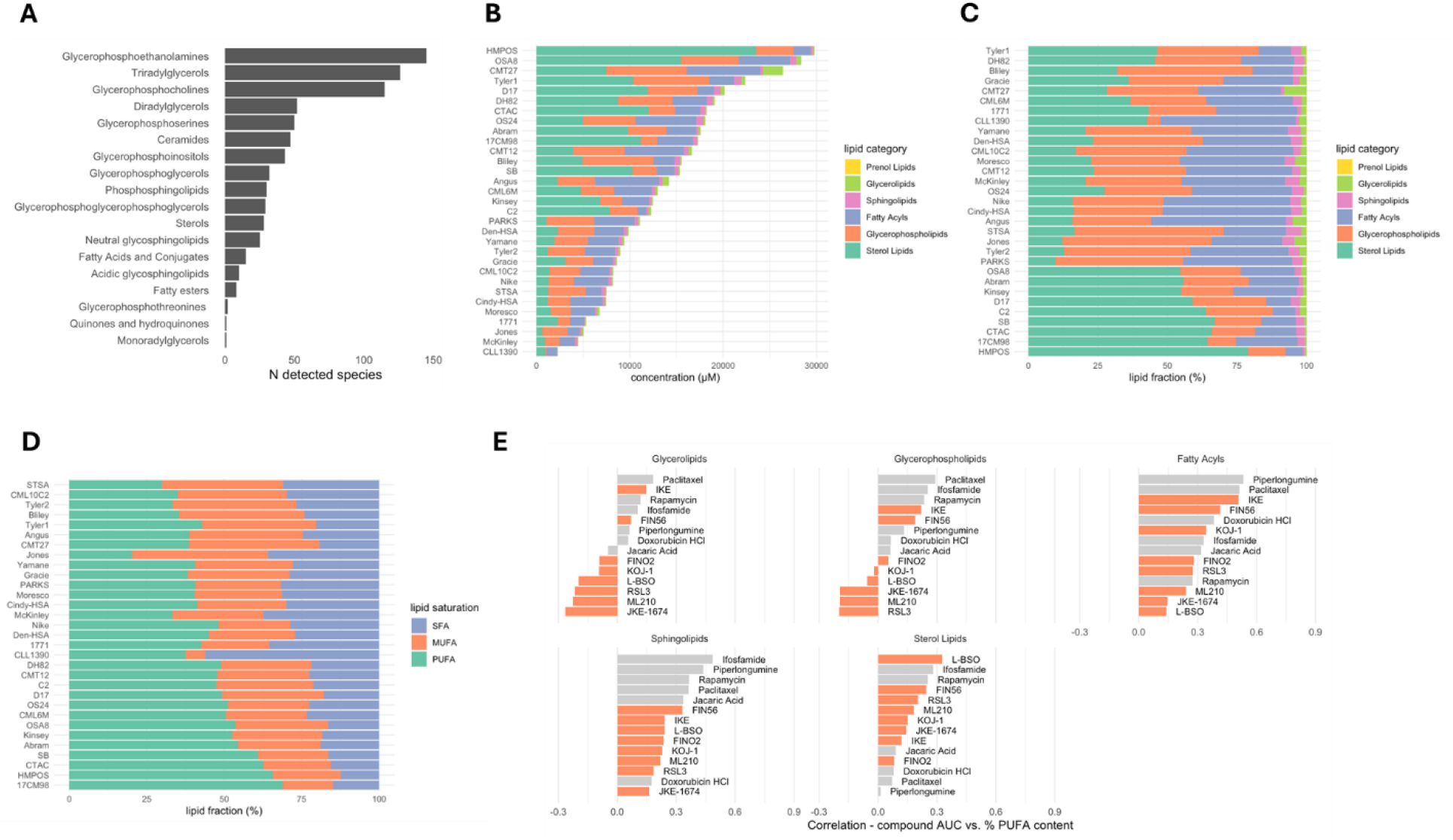
Canine cancer cell lines represent a wide range of lipidome profiles. (A) Detected lipid classes and number of individual species per class. (B) Lipid category composition of detected lipidome for each cell line; absolute concentrations. (C) Lipid category composition of detected lipidome for each cell line; relative values. (D) Distribution of lipid unsaturation for each cell line (SFA = saturated fatty acid, MUFA = monounsaturated fatty acid, PUFA = polyunsaturated fatty acid). (E) Pearson correlation between compound AUC values and relative PUFA content (%) for lipid categories. Negative correlation indicates that higher PUFA content corresponds to higher sensitivity (i.e., lower AUC).

While we did not observe strong correlations between total cellular PUFA content and sensitivity to ferroptosis inducers, higher PUFA content in glycerolipids and glycerophospholipids was associated with higher sensitivity to several ferroptosis inducers, particularly GPX4 inhibitors (Figure 4E). However, we anticipate that data from more cell lines are required to observe robust trends.

## Discussion

The ferroptosis pathway is regulated by several molecular mechanisms that have the propensity to vary greatly across species. For example, species-specific selenoproteomes and machinery for the biosynthesis of selenoproteins can lead to differences across species in the response to molecularly targeted ferroptosis inducers.^30,31^ Similarly, cellular lipid compositions, influenced by exposure to dietary lipids and expression of lipid synthesis and desaturating enzymes, are known to be divergent across species^32–37^ with potential impact on ferroptosis susceptibility. For these reasons, it is critical when exploring new model systems for ferroptosis translation to functionally test for sensitivity and assess whether key elements of the therapeutic rationale (e.g., heightened sensitivity of mesenchymal tumor types) are conserved. Here we have shown that canine cancers recapitulate patterns of ferroptosis sensitivity seen across human cancer cell lines^2^ and can serve as a valuable translational model. This work has also validated several cancer cell lines representing osteosarcoma and soft tissue sarcomas, which tend to be poorly represented among currently available human cancer cell line panels.^8^ Together, these results provide a strong foundation for undertaking further work to leverage canine comparative oncology its unique strengths to accelerate the generation of clinical proof of concept data for novel ferroptosis-inducing cancer therapeutics.

## Author contributions

V.S.V., J.K.E. and M.J.W. conceived of the project; P.C. performed experiments and coordinated the project; L.F. directed synthesis of compounds; Q.Z. performed cell line screening; D.V.L. directed cell line screening; D.H.T. enabled access to canine cancer cell lines and provided research input; G.X. oversaw lipidomic work; M.J.W. performed all data analysis, visualization and computational biology work; J.K.E., M.J.W. and V.S.V. wrote the paper.

## Acknowledgements

The authors thank the drug discovery and development shared resource (D3SR) high-throughput discovery sciences core. The D3SR is supported by the University of Colorado Anschutz Center for Drug Discovery, which was established with generous grant funding from The ALSAM Foundation. The D3SR is also supported in part by the University of Colorado Cancer Center, an NIH NCI designated center (P30-CA046934).

## Competing Interest Statement

V.S.V. is a co-founder and equity holder of Kojin Therapeutics. P.C., G.X., L.F., J.K.E., M.J.W., and V.S.V. are employees and equity holders of Kojin Therapeutics. L.F., J.K.E. and V.S.V. are inventors of patents related to ferroptosis.

---

**Table 1.**
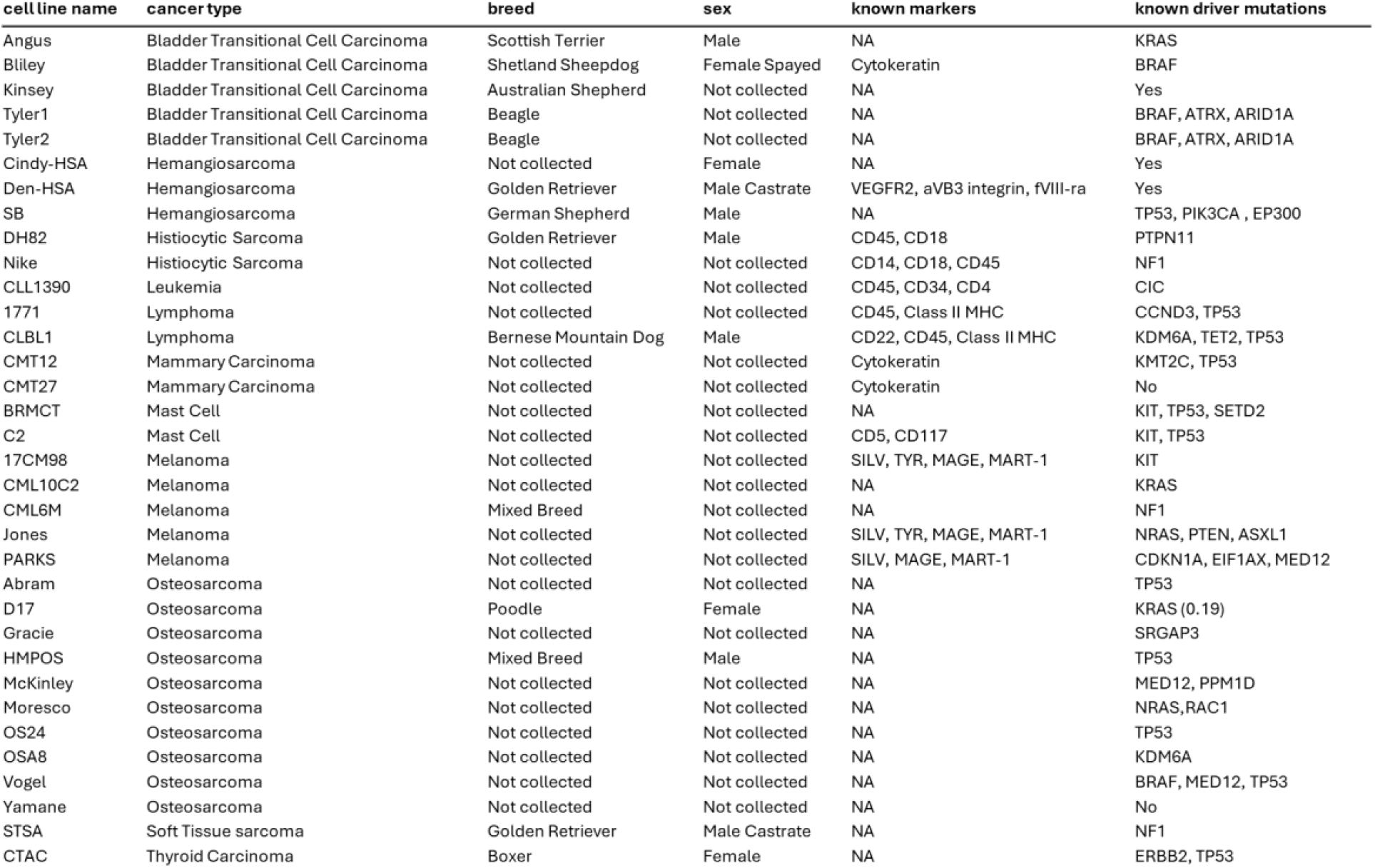
Summary of profiled canine cell lines.

**Supplementary Figure 1.**
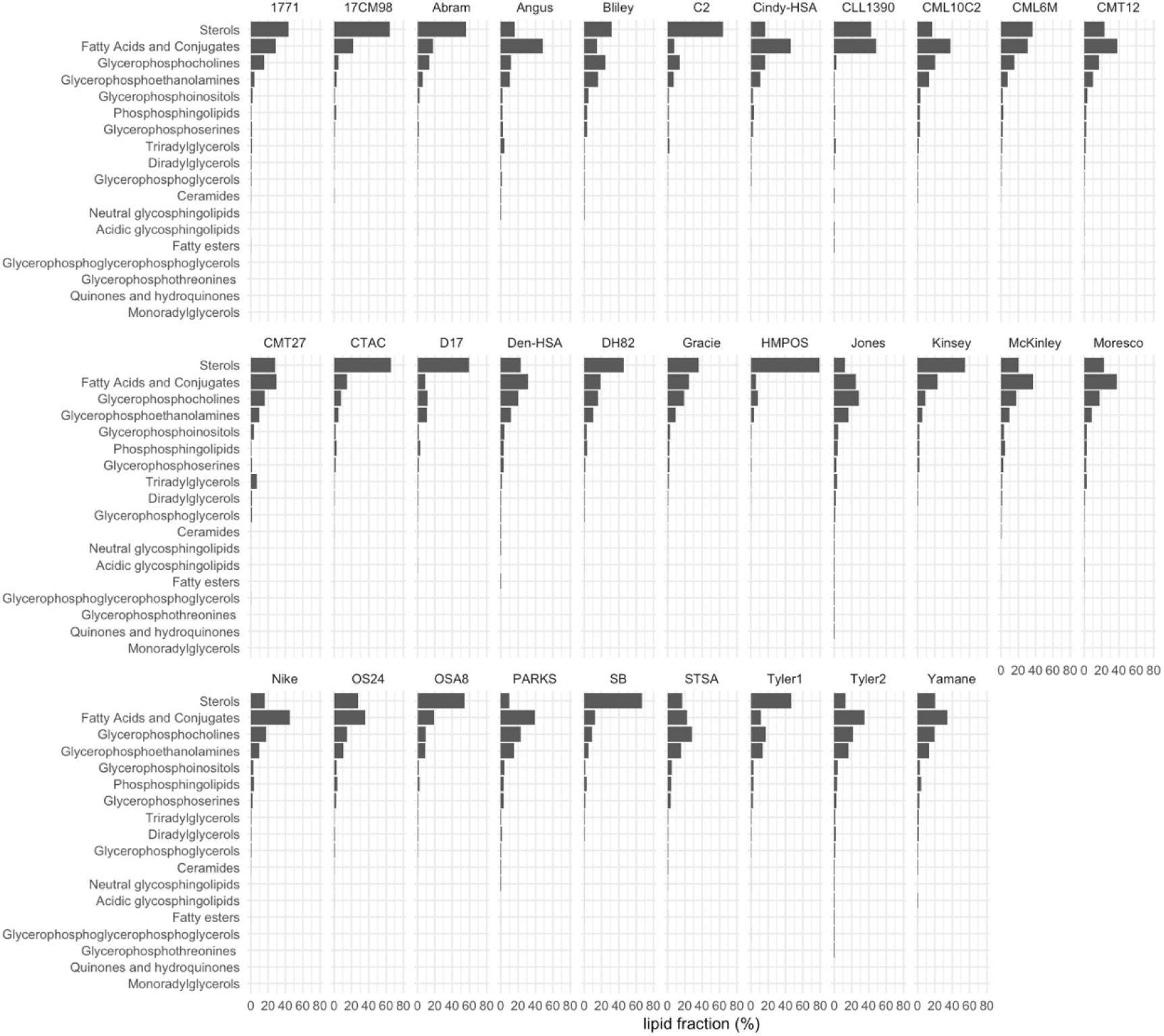
Lipid class composition for each cell line (semi-quantitative concentrations).

**Supplementary Figure 2.**
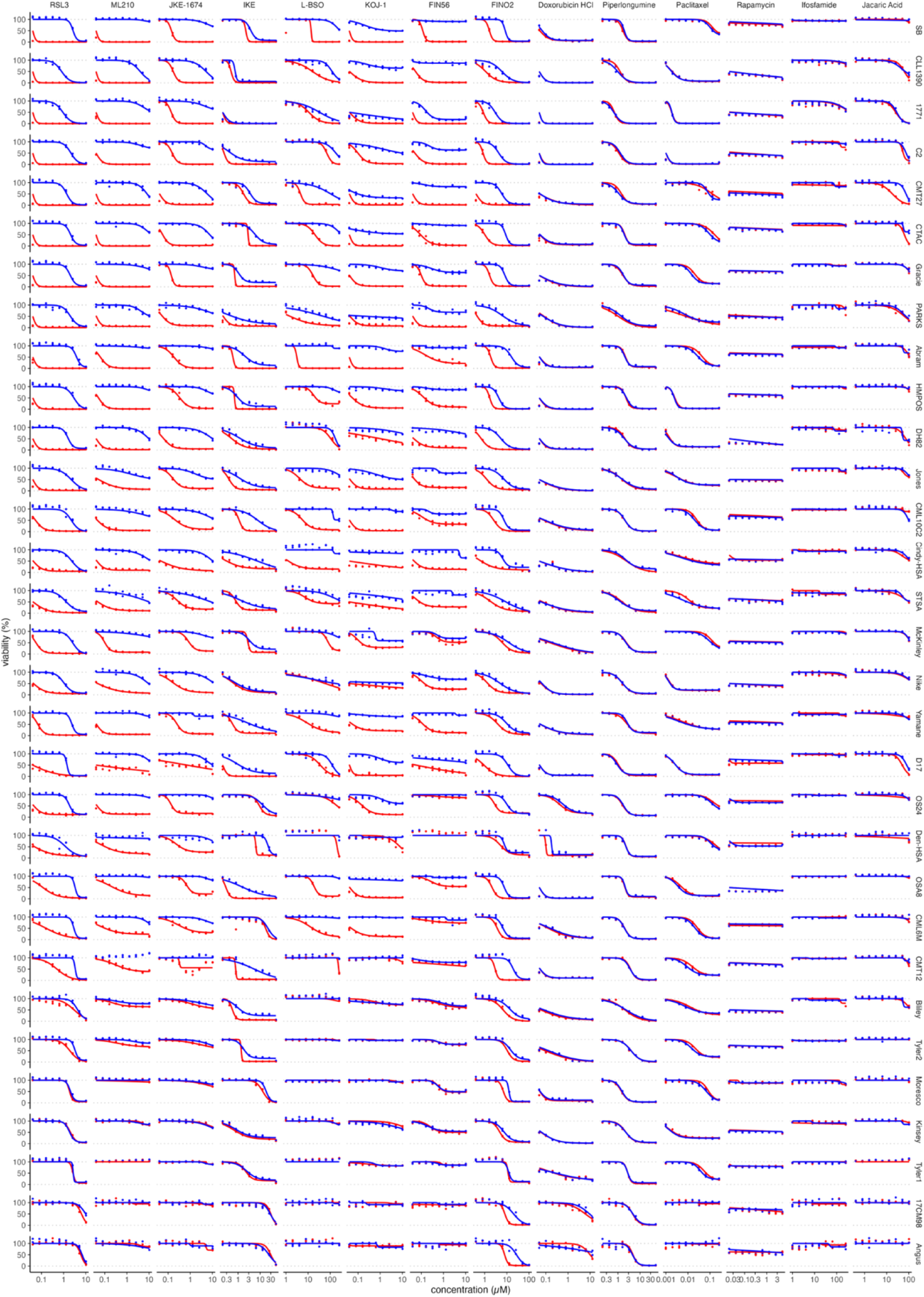
Dose–response curves for all compounds and cell lines.

